# PINOT: An Intuitive Resource for Integrating Protein-Protein Interactions

**DOI:** 10.1101/788000

**Authors:** JE Tomkins, R Ferrari, N Vavouraki, J Hardy, RC Lovering, PA Lewis, LJ McGuffin, C Manzoni

## Abstract

The past decade has seen the rise of omics data, for the understanding of biological systems in health and disease. This wealth of data includes protein-protein interaction (PPI) derived from both low and high-throughput assays, which is curated into multiple databases that capture the extent of available information from the peer-reviewed literature. Although these curation efforts are extremely useful, reliably downloading and integrating PPI data from the variety of available repositories is challenging and time consuming.

We here present a novel user-friendly web-resource called PINOT (Protein Interaction Network Online Tool; available at http://www.reading.ac.uk/bioinf/PINOT/PINOT_form.html) to optimise the collection and processing of PPI data from the IMEx consortium associated repositories (members and observers) and from WormBase for constructing, respectively, human and *C. elegans* PPI networks.

Users submit a query containing a list of proteins of interest for which PINOT will mine PPIs. PPI data is downloaded, merged, quality checked, and confidence scored based on the number of distinct methods and publications in which each interaction has been reported. Examples of PINOT applications are provided to highlight the performance, the ease of use and the potential applications of this tool.

PINOT is a tool that allows users to survey the literature, extracting PPI data for a list of proteins of interest. The comparison with analogous tools showed that PINOT was able to extract similar numbers of PPIs while incorporating a set of innovative features. PINOT processes both small and large queries, it downloads PPIs live through PSICQUIC and it applies quality control filters on the downloaded PPI annotations (i.e. removing the need of manual inspection by the user). PINOT provides the user with information on detection methods and publication history for each of the downloaded interaction data entry and provides results in a table format that can be easily further customised and/or directly uploaded in a network visualization software.

## Background

During the past two decades the use of omics data to understand biological systems has become an increasingly valued approach (1). This includes extensive efforts to detect protein-protein interactions (PPIs) on an almost proteome-wide scale (2, 3). The utility of such data has been greatly supported by primary database curation and the International Molecular Exchange (IMEx) Consortium, which promotes collaborative efforts in standardising and maintaining high quality data curation across the major molecular interaction data repositories (4). The primary databases, such as IntAct (5) and BioGRID (6), are rich data resources providing a comprehensive record of published PPI literature. PPI data are critical to describe connections among proteins, which in turn supports both inference of new functions for proteins (based on the guilt by association principle (7)) and visualization of protein connectivity via shared interactors. This support shedding light on communal pathways involving proteins of interest (8–10). Additionally, literature extracted PPI data can support the prioritization of interactions from high-throughput experiments (which generate large lists of potential PPI hits), assisting the selection of candidates for further analysis/validation (11).

However, the process of collating PPI data from multiple sources is currently hampered by the fact that no single data source encompasses the full extent of PPIs reported in the literature, requiring users to merge (partial) information mined from different primary databases. Furthermore, merging such data is not straightforward due to inconsistencies in data format and differences in data curation across the PPI databases (IMEx members vs non-members).

To optimize the use of PPI data from the public domain, we developed a user-friendly tool that assists PPI data extraction and processing: the Protein Interaction Network Online Tool (PINOT). This tool represents the development (and automation) of our previous PPI analysis framework (i.e. weighted protein-protein interaction network analysis – WPPINA) (9, 11–15). Through PINOT, PPI data is downloaded directly (i.e. downloaded “live” at the time of the query) from seven databases using the Proteomics Standard Initiative Common Query Interface (PSICQUIC) and integrated to ensure a wide coverage of the PPIs available from these repositories (16). These data are scored through a simple and transparent procedure based on ‘method detection’ and ‘publication records’ and allows the user to further apply customized confidence thresholds. PINOT is fully automated and available online as an open access resource. Output data are provided as a summary table (directly online or emailed to the user), which summarizes the most comprehensive current knowledge of the PPI landscape for the protein(s)-of-interest submitted in the query list. Of note, the R scripts which underlie PINOT can be freely downloaded from the help-page.

## Methods

### Protein Interaction Network Online Tool (PINOT)

PINOT can be run automatically at http://www.reading.ac.uk/bioinf/PINOT/PINOT_form.html (hereafter referred to as “webtool”). A choice of parameters is integrated by default as explained further below and in Supplementary Materials (S1). Alternatively, R scripts can be downloaded from the help-page (hereafter referred to as “standalone tool”, since parameters can be modified as *per* user choice).

A list of proteins of interest (seeds) can be queried to identify their literature-reported interactors that have been curated into PPI databases (Figure 1).

**FIGURE 1.**
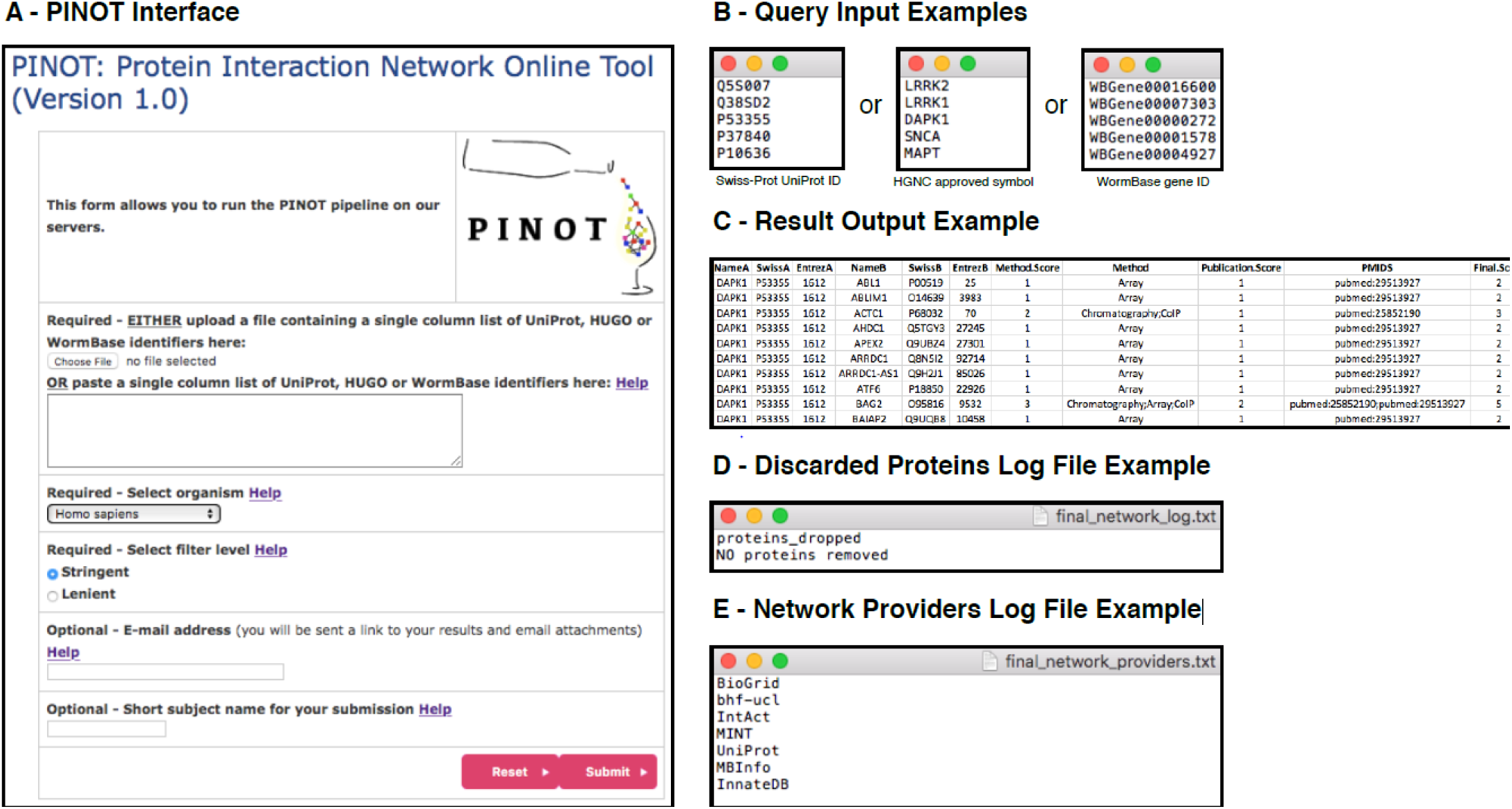
PINOT user interface. A. Screenshot of the PINOT webpage, B. Examples of the text file to be uploaded or list to be populated into the text box of query seeds (i.e. proteins for which protein interactors will be extracted from primary databases that manually curate the literature), C. Example result output file from PINOT, containing the extracted and processed PPI data (only the file’s header is reported as an example), D. Example of the discarded proteins log file from PINOT, a text file reporting all the seeds for which interactions are not returned to the user, and E. Example of the network providers log file from PINOT containing a list of active databases that were utilised for downloading PPI data.

For *Homo sapiens* (taxonomy ID: 9606) the seed identifiers submitted into the query field must be in an approved HUGO Gene Nomenclature Committee (HGNC) gene symbol or valid Swiss-Prot UniProt ID format. Upon query submission, PPI data are extracted directly (*via* API: Shannon, P. (2018) PSICQUIC R package, DOI: 10.18129/B9.bioc.PSICQUIC (17)) from seven primary databases, all of which directly annotate PPI data from peer-reviewed literature: bhf-ucl, BioGRID (6), InnateDB (18), IntAct (5), MBInfo (https://www.mechanobio.info), MINT (19) and UniProt (20). The downloaded protein interaction data are then parsed, merged, filtered and scored (Figure 2) automatically by PINOT. Detailed description of the PINOT pipeline can be found in the supplementary materials (S1). The user can select to run PINOT with lenient or stringent filter parameters. The output of PINOT (Figure 1C-E) consists of: i) a network file (final_network.txt), which is a tab-spaced text file containing the processed PPI data in relation to the seeds in the initial query list; ii) a log file (final_network_log.txt) reporting proteins that have been discarded from the initial query list, and; iii) a log file (final_network_providers.txt) indicating the PPI databases used by the API at download. The output dataset is available for download and/or emailed to the user.

**FIGURE 2.**
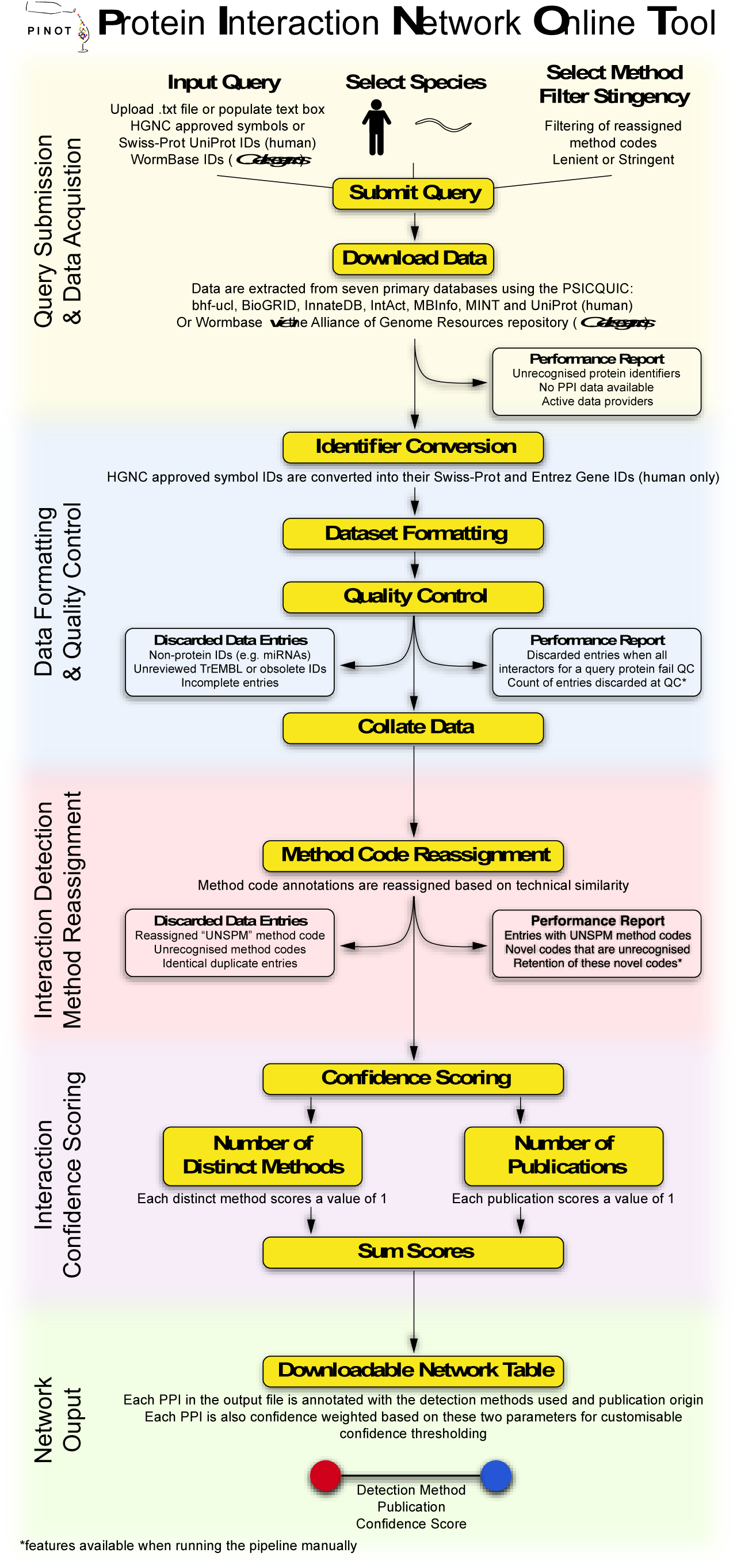
PINOT pipeline. A stepwise insight into the process which underlies the PINOT pipeline. Performance reports (green boxes) are generated and data are discarded (red boxes) at numerous stages within the pipeline to ensure high quality and transparent data processing.

For *Caenorhabditis elegans* (taxonomy ID: 6239) the seed identifiers must be in an approved WormBase gene ID (21) format, “WBGene” followed by 8 numerical digits. Upon submission PPI data are downloaded from an internal network stored within PINOT and created (following similar criteria applied for the human PPIs – details in S1) based on the WormBase PPI catalogue (Alliance_molecular_interactions.tar file downloaded from the Alliance of Genome Resources on 15th April 2019). The user can apply stringent or lenient filtering options. The output of PINOT for a *C. elegans* query consists of: i) a network file (final_network.txt), which is a tab-spaced text file containing the processed PPIs for the seeds in the initial query list; and ii) a log file (final_network_log.txt) reporting proteins that have been discarded from the initial query list.

### Software

The PINOT pipeline is coded in R and runs on a Linux server at the University of Reading, with java servlets processing user’s submissions *via* the web interface.

### PINOT quality control

We have tested the PINOT pipeline using multiple input query lists structured as follows: i) small input lists = 6 sets of 1 to 5 proteins, selected randomly or in association with typical processes suspected to be functionally relevant for Parkinson’s Disease (PD); and ii) large input list = 941 proteins, the mitochondrial proteome as reported by MitoCarta2.0 (22).

PINOT was compared to two other related online tools. For this analysis, searching parameters were selected (where possible) to maximize the extraction of protein interactions: the Human Integrated Protein-Protein Interaction Reference (HIPPIE) was used with confidence score = 0 and no filters on confidence level, interaction type or tissue expression; and the Molecular Interaction Search Tool (MIST) was used with no filtering rank to download only protein protein interactions. Importantly and of note, files from HIPPIE and MIST required manual parsing after download to remove entries that were associated to no PMID and/or no conversion method code (incomplete entries). Data were downloaded on 18th September 2019 (*H. sapiens*) and on 24^th^ September 2019 (*C. elegans*).

## Results

PINOT is a webtool that takes a list of proteins/genes (seeds) as input and returns a table containing a comprehensive list of PPIs – published in peer-reviewed literature – centred upon the seeds. This table consists of a variable number of rows and 11 columns (Figure 1C and 3C). Each row represents a binary interaction between one of the seeds (interactor A) and one of its specific protein interactors (interactor B). The 11 columns contain: the gene name, the Swiss-Prot protein ID and the Entrez gene ID for interactor A and B (“NameA”, “SwissA”, “EntrezA”, “NameB”, “SwissB”, “EntrezB”); the number and type of different methods through which the interaction has been identified (“Method.Score”, “Method”); and the number of different publications reporting the interaction and the corresponding PubMed IDs (“Publication.Score”, “PMIDS”). The final column (“Final.Score”) contains a confidence score based on the number of different methods + the number of different publications reporting the interaction. PPIs with a final score of 2 are reported in literature by 1 publication and detected by 1 technique; these PPIs are considered “suggestive” (but are clearly not “replicated”). They might be either: i) false positives, or ii) true novel interactions that have not yet been replicated in additional studies. A final score >2 suggests a degree of replication that can be either or both: multiple publications reporting the PPI and multiple techniques used to detect the interaction. It is not possible to obtain a final score <2 since every PPI annotation – to be retained in PINOT – has to be supported by at least 1 interaction detection method and 1 PMID; if this condition is not met, the PPI is discarded by PINOT and not shown in the output file.

The PINOT output can be imported into Cytoscape (23) for network visualization by selecting the “NameA” and “NameB” columns as source and target nodes, respectively.

### PINOT: Example of application

In Figure 3 PINOT has been used to download PPIs for a limited selection of human protein products of genes mutated in familial PD: ATP13A2, FBXO7, GBA, PINK1, SMPD1 and VPS35 (seeds). PINOT quickly retrieved a table containing 327 interactions from peer-reviewed literature (with associated PMIDs) thus supporting and simplifying otherwise time-consuming classical literature mining. The PINOT output was imported into Cytoscape and PPIs were visualized in a network (“NameA” = source and “NameB” = target), the seeds were highlighted in dark-red and the edges (interactions between each protein) were coded based on the “Final.Score” field, thus highlighting the confidence (number of methods + number of publications) of the interaction. Since we were interested in interactors that were common to the seeds – and not in “private” interactors of just one seed – the network was filtered retaining only the nodes (interactors) that bridged two or more seeds. The obtained core-network revealed that among the common interactors of the seeds (PD proteins) there were 2 proteins (SNCA and PRKN), which are products of 2 additional genes known for being mutated in familial PD. Thus, the analysis pointed towards the involvement of SNCA and PRKN in PD even if they were initially excluded from the list of seeds. Additionally, topological analysis (based on the number and strength of the edges) suggested the core network could be subdivided into 2 distinct clusters respectively including PINK1, FBXO7 and the newly identified PRKN and SNCA in the first cluster, while ATP13A2, VPS35 and SMPD1 were more closely associated in the second cluster, with GBA a bridge seed between the 2 clusters. This observation suggested a dichotomy, based on the protein interactomes, of the seeds included in the initial input list. Based on the guilt-by-association principle we hypothesised that the proteins contributing to these clusters could be associated with different cellular functions and components. We therefore performed functional enrichment analysis (based on Gene Ontology (GO) Cellular Component (CC) annotations) using g:Profiler (24) revealing that indeed, clusters 1 and 2 are associated with mitochondria and vacuoles/lysosomes/endosomes, respectively.

**Figure 3.**
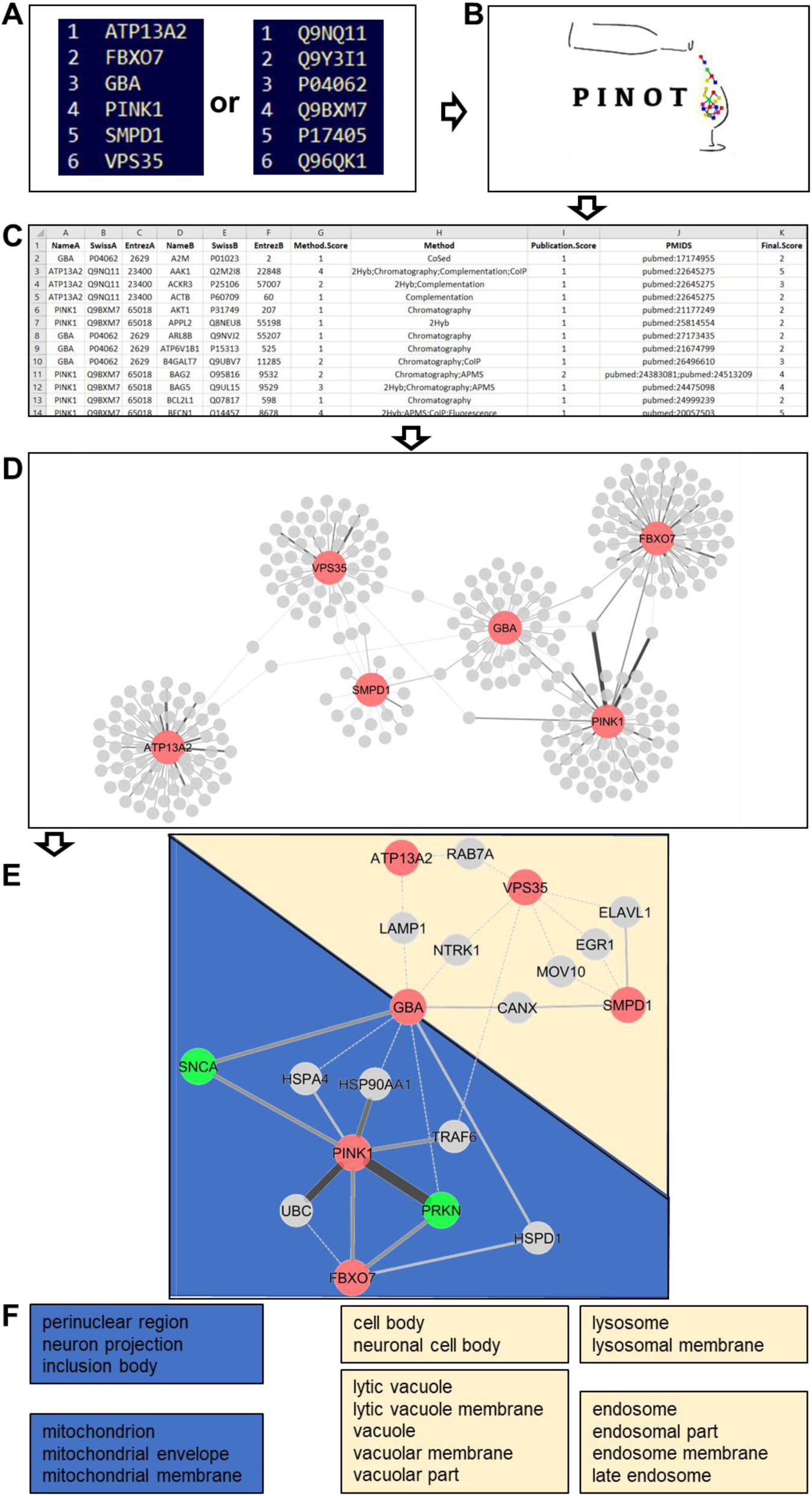
PINOT: An example application. A stepwise insight into the potential use of PINOT. 1. A submission list is created as a text file using gene names as per HGNC approved symbols or Swiss-Prot IDs; the submission list can be uploaded as file or pasted into the PINOT interface. 2. PINOT downloads from PSICQUIC the human PPIs (in this example, stringent filters applied) 3. PPIs are provided back to the user *via* email or from the webpage; results are in a parsable file that can be opened by a text reader application and imported into Microsoft Excel. 4. The interactions can be visualized in a network format by opening the PINOT output through Cytoscape. Connections between nodes (edges) are coded with increased line width based on the final score that interaction was assigned by PINOT. The wider the edge – the more confident PINOT is about the interactions. 5. The interactions can be further processed according to the user’s research question, in this case, only interactors that are communal to at least 2 of the initial query proteins have been retained, generating a core network (in dark-red the initial seeds; in bright-green the identified common interactors that are proteins mutated in PD). Based on the network topology the seeds and their interactors can be visually clustered into group 1 (depicted in gold) and group 2 (depicted in blue). 6. Specific functional enrichment (GO CC terms) for groups 1 and 2 after filtering out the less represented terms. Analyses performed on the 22^nd^ August 2019.

### H. sapiens – PINOT performance

The performance of PINOT was compared to that of alternative resources for both small and large numbers of seeds. Regarding the former, five different small seed lists were used as input for PPI query in HIPPIE (25) and MIST (26), two alternative online and freely available resources. It should be noted that, despite apparent similarities, each of these tools has been developed differently. All three resources (PINOT, HIPPIE and MIST) have distinguishing features for addressing different research questions (Table 1). The results of the different queries have been compared, evaluating the total number of interactors provided in the output (Figure 4A).

**Table 1.**
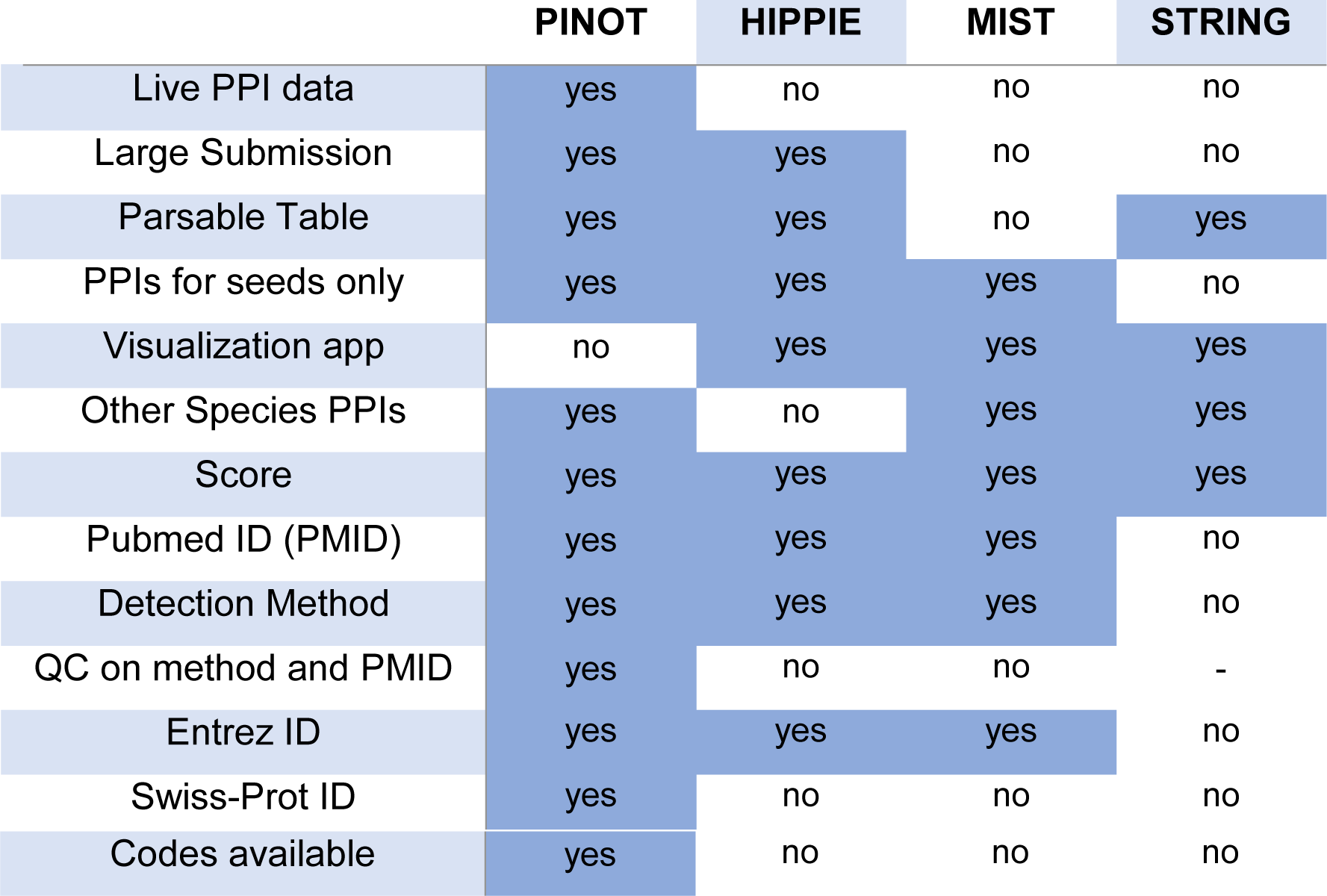
Comparison of features available within the PINOT, HIPPIE, MIST and STRING resources.

|  | PINOT | HIPPIE | MIST | STRING |
| --- | --- | --- | --- | --- |
| Live PPI data | yes | no | no | no |
| Large Submission | yes | yes | no | no |
| Parsable Table | yes | yes | no | yes |
| PPIs for seeds only | yes | yes | yes | no |
| Visualization app | no | yes | yes | yes |
| Other Species PPIs | yes | no | yes | yes |
| Score | yes | yes | yes | yes |
| Pubmed ID (PMID) | yes | yes | yes | no |
| Detection Method | yes | yes | yes | no |
| QC on method and PMID | yes | no | no | - |
| Entrez ID | yes | yes | yes | no |
| Swiss-Prot ID | yes | no | no | no |
| Codes available | yes | no | no | no |

**Figure 4.**
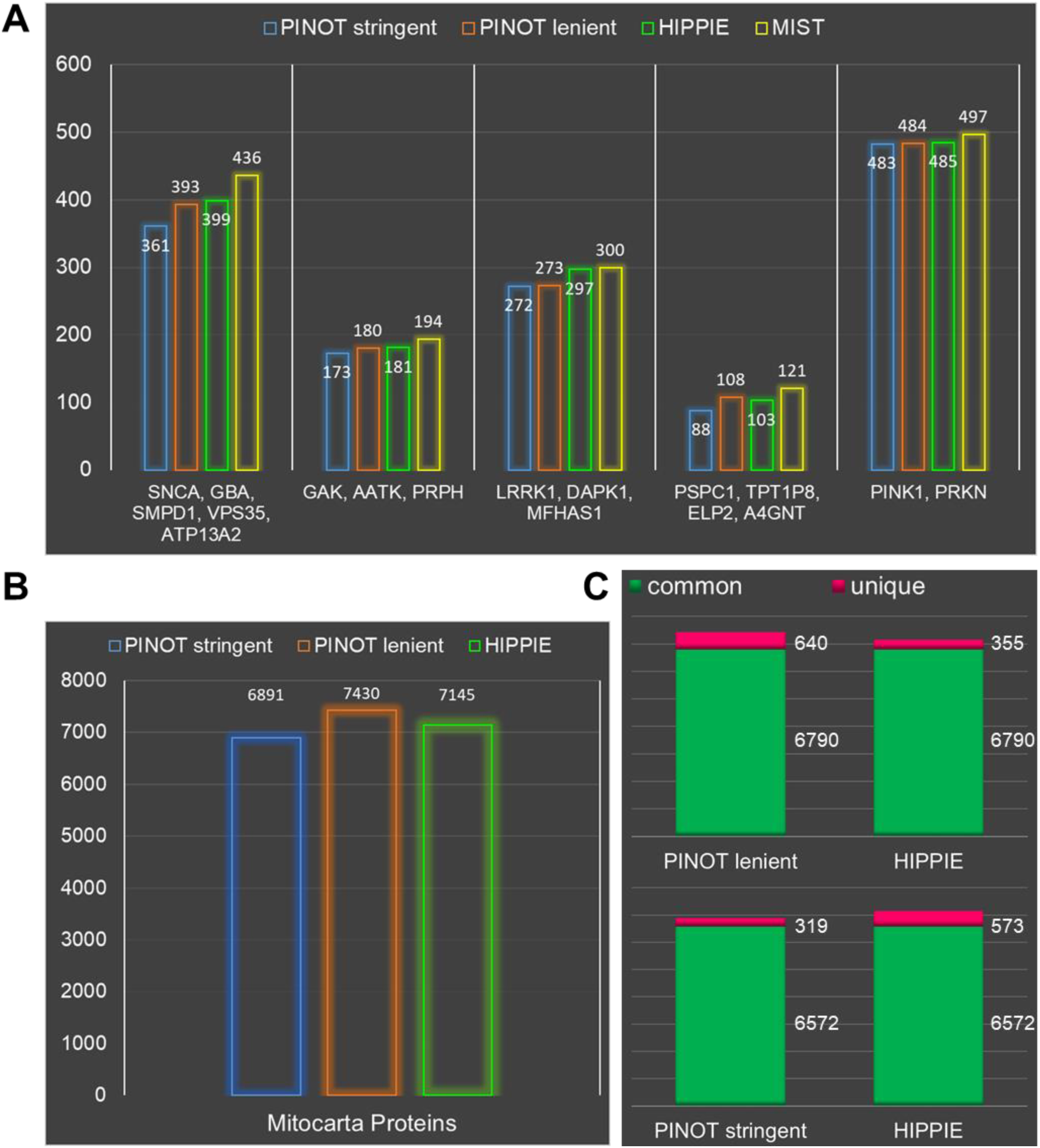
PINOT: Performance & Sensitivity. A. PINOT performance was evaluated by counting the number of interactors retrieved (gene names) upon submission of the reported query lists to PINOT (with stringent and lenient filtering), HIPPIE and MIST (on 18th September 2019). The databases were set to retrieve the maximum number of interactions (by removing all possible filters). The HIPPIE and MIST outputs were manually cleaned to remove interactions with i) no interaction detection method; no PMID; iii) multiple Entrez IDs. The number of retained interactions retrieved is reported on top of each bar. B. PINOT (with stringent and lenient filtering) and HIPPIE were queried to retrieve PPIs for a seed list of 941 protein from Mitocarta 2.0. C. Comparison between PINOT and HIPPIE showing that the vast majority of interactors (Entrez IDs) downloaded by the two tools was identical: 6790 common interactors for PINOT lenient (640 unique interactors) vs HIPPIE (355 unique interactors); 6572 common interactors for PINOT stringent (319 unique interactors) vs HIPPIE (573 unique interactors).

PINOT, HIPPIE and MIST retrieved a similar number of PPIs. PINOT with stringent filtering applied, was always extracting fewer interactions; this is an expected outcome since this filter option is built with the purpose of retaining only annotations that have survived stringent screening, largely based on completeness of curated data entries. The large input list was compared in PINOT and HIPPIE, the only two webtools that allowed for easy processing of more than 900 seeds within the submission list. In fact, MIST submission needed to be divided into multiple small lists to allow the browser to properly process the query. Additionally, the downloaded table(s) were not parsable (in an automated fashion), thus making MIST (the version available at the time of the query) counterintuitive for the processing of large input lists. The number of retrieved interactors was slightly higher for HIPPIE in comparison with PINOT when the stringent QC filter was applied; while PINOT with lenient filtering retrieved more interactions than HIPPIE (Figure 4B). Additionally, the vast majority of downloaded interactions were similar from using the two resources, suggesting that PINOT is able to confidently extract specific interations from literature (Figure 4C).

### C. elegans – PINOT performance

The performance of PINOT for querying *C. elegans* PPI data was tested alongside the *C. elegans* query option in MIST, assessing interaction networks of different dimensions (Figure 5). The data acquisition strategy underlying these two resources differs slightly, PINOT extracts data from the latest release of WormBase molecular interaction data, whereas MIST utilises data from numerous sources, including WormBase, BioGRID and IMEx associated repositories.

**Figure 5.**
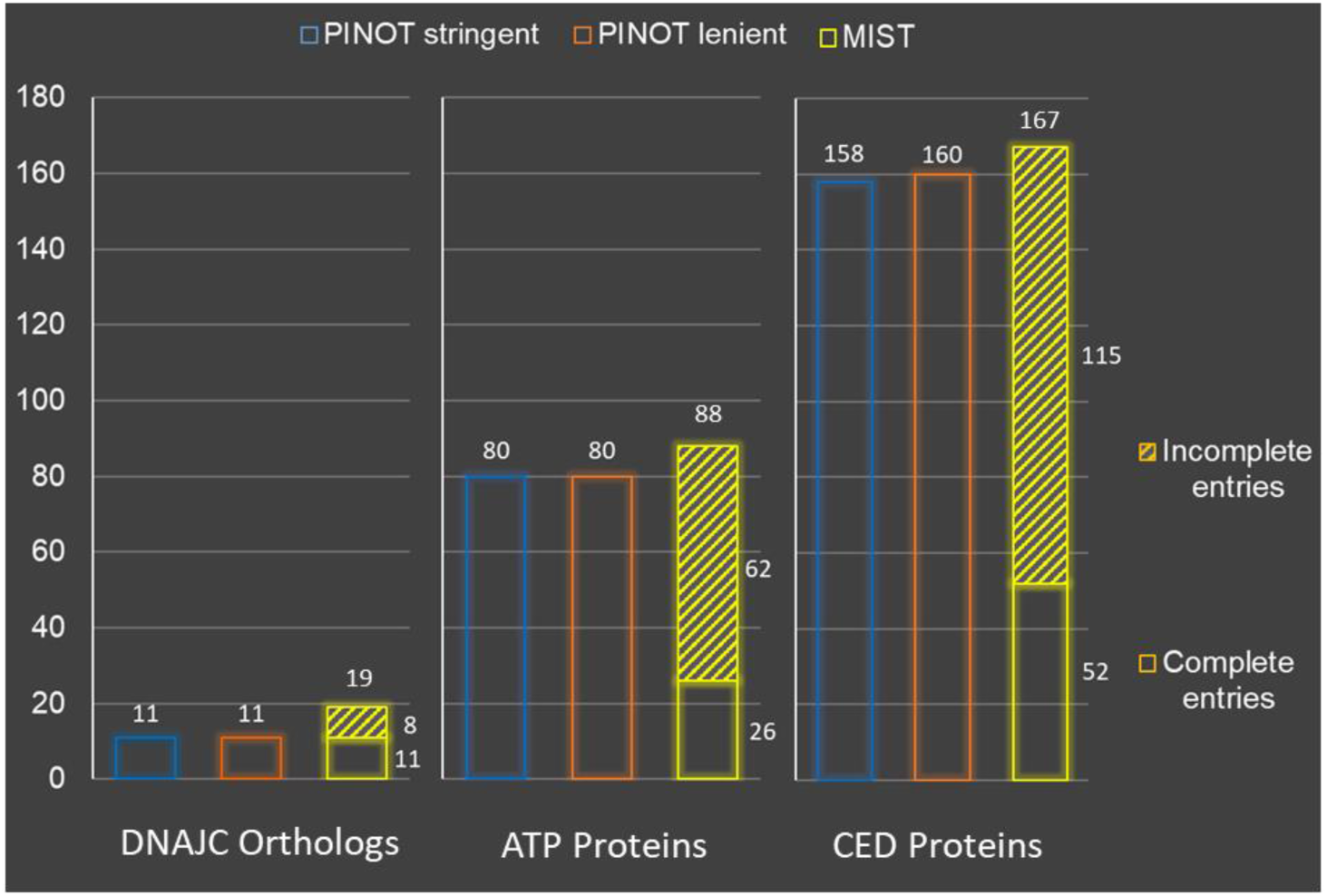
PINOT and MIST performance comparison for *C. elegans* PPI data. The performance of PINOT (with stringent and lenient filter options) and MIST was assessed in terms of the number of PPI data entries extracted upon querying different protein lists (on 24th September 2019). The output dataset was evaluated in relation to the number of complete and incomplete (lacking interaction detection method and/or PubMed ID annotations) data entries extracted. The query lists were PD-associated DNAJC orthologs: DNJ-14, DNJ-25, DNJ-27, Y73B6BL.12, K07F5.16, RME-8 and GAKH-1; ATP proteins: ATP- 1, ATP-2, ATP-3, ATP-4, ATP-5 and ATP-6; and CED proteins: CED-1, CED-2, CED-3, CED- 4, CED-5, CED-6, CED-7, CED-8, CED-9, CED-10, CED-11, CED-12 and CED-13. The input format used for PINOT was the WormBase gene ID, the common gene name (as listed here) was used for MIST querying and no filter by rank parameter was set.

Similarly to comparisons across the human PPI query capacity, PINOT and MIST performed comparably in terms of the number of PPI data entries extracted. More specifically and as previously described with human data, PINOT extracting slightly fewer across these test query cases. However, upon assessing the completeness of these extracted data entries, in terms of interaction detection method and/or PMID annotations, there was a striking difference in performance. Since the PINOT pipeline focusses particular emphasis on the QC of data, all data entries within the output dataset were complete, whereas incomplete data entries persisted in the MIST output dataset thus requiring manual inspection. In the more abundant PPI data pools, for example when querying the ATP and CED *C. elegans* proteins, incomplete data entries accounted for the majority of the output dataset in MIST.

## Discussion

PINOT can be used as a tool to quickly and effectively survey the literature and download the most up-to-date PPI data available for a given set of proteins/genes of interest. This is particularly useful to assist anyone attempting to mine overwhelming abundant literature targeting certain proteins/genes, in relation identifying reported PPIs.

The PPI data downloaded through PINOT can be used as a reference list (from literature) for experimental PPI data resulting from high-throughput experiments (protein microarrays; yeast 2 hybrid screens, etc) helping in prioritisation of experimental results for validation. PINOT can also be useful to evaluate interactors of different proteins/genes of interest within an input seed list simultaneously. The analysis of the combined interactomes of such seeds can reveal the existence of communal interactors, can provide a base to cluster the seeds into groups and can support further functional analysis to better characterize the functional landscape of seeds of interest.

Alternative tools that appear to be similar to PINOT are HIPPIE and MIST. STRING (27) is a conceptually different tool; it does not report ‘interaction detection methods’ nor ‘Publication IDs’ for PPIs which are crucial pieces of information for the evaluation and interpretation of PPI data. Additionally, the reported interactions are not focused only on the proteins in the input list; interactions of interactors are also reported, thus making it difficult to parse the output table. HIPPIE implements a tailored confidence score for different methodological approaches; MIST provides a valuable resource for users interested in mapping PPIs across species (i.e. interologs); PINOT focusses on high quality PPI data output by implementing multiple QC steps to remove problematic or non-univocal annotations. PINOT performance was comparable to that of HIPPIE and MIST both in terms of number and identity of downloaded interactions. However, there are some unique features of PINOT that are not, at the moment, integrated within the other databases. Human PPIs in PINOT are directly downloaded from PSICQUIC at every query submission. In contrast, PPIs in HIPPIE and MIST are recovered from a central built-in repository within the servers. This difference is clearly demonstrated by searching for interactors of LRRK2, where (at the time of analysis) 1 high-throughput publication was updated in PSICQUIC, while both HIPPIE and MIST did not contain this full annotation yet (Figure 6).

**Figure 6.**
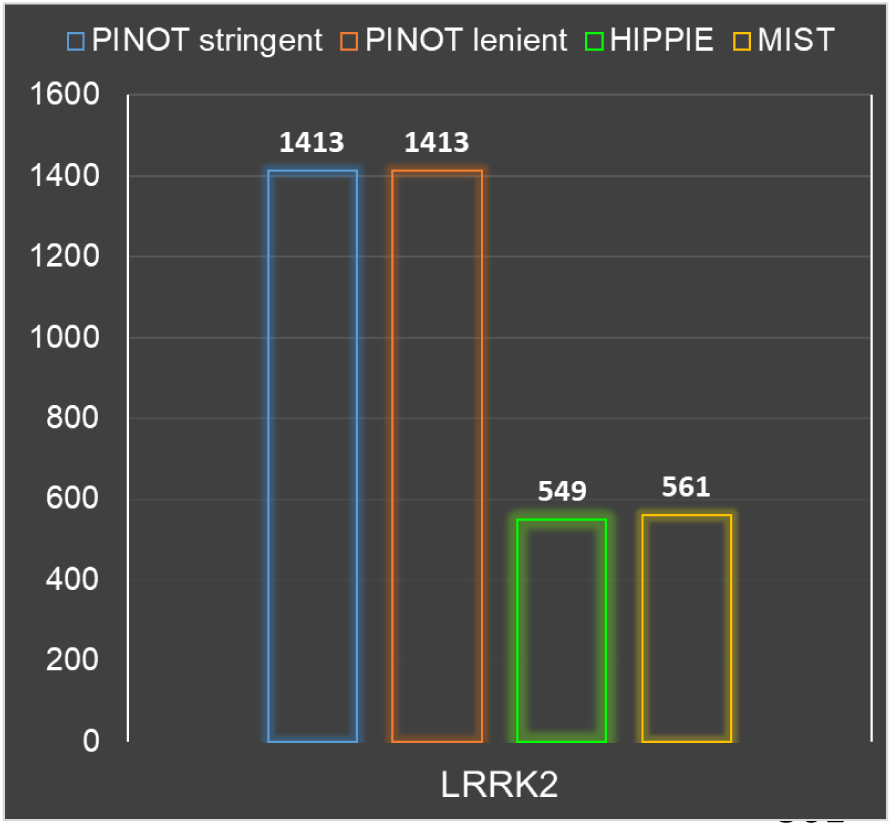
LRRK2 interactome. PINOT performance was evaluated by counting the number of interactors retrieved (gene names) for LRRK2 by using PINOT (with stringent and lenient filtering), HIPPIE and MIST. The databases have been set to retrieve the maximum number of interactions (by removing all possible filters). HIPPIE and MIST output were manually cleaned to remove interactions with i) no interaction detection method; ii) no PMID; iii) multiple Entrez IDs. The number of the surviving interaction retrieved is reported on top of each bar (18^th^ September 2019).

PINOT has access to the most up-to-date interactions that could be retrieved at a given time from PSICQUIC (however, it has to be considered that each database is responsible for updating their PSICQUIC service and therefore discrepancies might exist with the central database).

PINOT implements QC filtering which involves discarding PPI data entries that are curated without a PMID and/or the interaction detection method annotation. Therefore the output file from PINOT does not require any further QC by the user, while lists from MIST and HIPPIE require manual parsing and inspection before analysis to remove incomplete data entries through a time consuming, post-hoc processing procedure. Another distinctive feature of the PINOT pipeline is the implementation of a unique interaction detection method conversion step. During this step, the interaction detection method annotation for each downloaded interaction data entry is converted based on a conversion table (S2) that is available for download from the PINOT web-portal. During this conversion, technically similar methods are grouped together. For example: “Two Hybrid – MI:0018”, “Two Hybrid Array – MI:0397” and “Two Hybrid Pooling Approach – MI:0398” are grouped together into the “Two Hybrid” method category. This step of ‘method clustering and reassigment’ is critical to assess the actual number of distinct methods used to describe a particular interaction and to dilute the bias caused in the event of the same technique being annotated under slightly different method codes in different PPI databases.

Interaction scores are provided in different formats for the three tools. HIPPIE incorporates a filtering system based on a confidence score between 0 and 1 that can be set either before or after the analysis. This is a complex scoring system, which takes into consideration multiple parameters, such as the number of publications that report a specific interaction and a semi-computational quality score based on the experimental approach (for example, imaging techniques would score less than direct interactions etc.) (28). MIST similarly has an option for filtering interactions pre- or post-analysis; however, this is based on fixed ranking values defined as low, medium (interaction supported by other species), or high (supported by multiple experimental methods and/or reported in multiple publications). In the case of PINOT, two different scores are provided: the interaction detection method score (MS) reports the number of different methods used (after conversion), while the publication score (PS) counts the number of different publications which report the interaction. Finally, *H. sapiens* PINOT coding scripts are fully available for download. They are coded in R to make them accessible to a large research audience; additionally a read me text file helps customization of the scripts according to the users’ needs. Some of the divergent features across PINOT, HIPPIE, MIST and STRING are reported in Table 1.

## List of Abbreviations

HIPPIE: Human Integrated Protein-Protein Interaction Reference
MIST: Molecular Interaction Search Tool
MS: method score
PPI: protein protein interaction
PD: Parkinson’s Disease
PINOT: protein interaction network online tool
PMID: Pubmed ID
PS: publication score
PSICQUIC: Proteomics Standard Initiative Common Query Interface
QC: Quality Control
WPPINA: weighted protein-protein interaction network analysis.

## Availability of data

This resource is available as a fully automated web-server at: http://www.reading.ac.uk/bioinf/PINOT/PINOT_form.html; R scripts, which underlie this bioinformatics pipeline, are free for download at the help-page.

## Author’s Contribution

TEJ, MC and FR, elaborated the pipeline and wrote the manuscript; TEJ, MC and VN wrote the scripts and tested the pipeline, MJL implemented the pipeline for the website, HJ, LCR, LAP offered critical advice for the implementation of the pipeline, critically reviewed the manuscript and obtained financial support.

## Acknowledgements

We thank WormBase for providing the list of PPIs used for the generation of the internal *C. elegans* network used when *C. elegans* genes/proteins are queried by PINOT users.

## Supplementary Files

**S1** = supplementary materials and methods

**S2** = supplementary file 1: ‘interaction detection method conversion’

