## Supplementary S1 for "PINOT: An Intuitive Resource for Integrating Protein-Protein Interactions"

### *Homo sapiens (taxonomy ID: 9606)*

#### *Query Submission and Data Acquisition*

A file is created for each seed protein from the input query list, which consists of interaction data entries captured from all the interrogated primary databases. In this file, each row reports a single curated interaction consisting of interaction partner A and interaction partner B where one of the two partners (A or B) is the seed. A report on the active data providers at the time of the query is generated and provided to the user. This allows the user to determine the origin of the output data and identify whether any of the databases were offline at the time of query. If a query protein ID is not recognised (i.e. the protein identifier is not provided in the approved HUGO Gene Nomenclature Committee (HGNC) or Swiss-Prot formats a warning message is generated.

#### *Data Processing: Data Formatting, Quality Control (QC) & Filtering*

To integrate information captured from different primary databases, each file generated from the initial query is parsed and QCed.

1. All protein identifiers are converted into their Swiss-Prot ID, Entrez ID and HGNC symbol using a dictionary list obtained from processing the <hgnc_complete_set> file downloaded from HGNC: (https://www.genenames.org/download/statistics-and-files/) on August 2019. Briefly, genes and proteins that are univocally coded by Swiss-Prot and Entrez are kept in the dictionary while non-univocal entries and genes with no associated protein ID are dropped.

2. Interactors annotated with non-protein IDs (e.g. chemical or miRNA interactors) and interactors that are annotated with non-reviewed TrEMBL or obsolete protein IDs are discarded from further data processing. Proteins in the input query list (seeds) that do not have any interactors surviving this first QC step are removed and the information captured in a log file. Note that a protein from the initial query list might be associated in the primary databases with an obsolete protein ID, in this case the protein is discarded (see above). The removal of obsolete protein IDs

cannot be modified while using PINOT in its automated version, however this can be overcome while using the custom tool by retrieving and correcting the removed file from the LogFiles folder.

3. Each file is then consistently formatted in respect to the position of specific information in the designated columns (i.e. interactors A and B are ordered so that the initially queried protein is always in the A position).

4. Data undergoes further QC checks based on how the publication reporting each binary interaction is annotated within the primary database. Specifically, interaction entries are discarded if they are: i) annotated without a complete PMID, or ii) annotated with multiple PMIDs. Proteins in the input query list (seeds) that do not have any interactors surviving the publication QC step are removed and annotated in a log file.

5. To overcome the annotation bias for which two different primary databases may have annotated the same interaction (from the same publication) with (slightly) different interaction detection method codes (this would generate a bias at the scoring step), the interaction detection method is reassigned based on method similarity. Methods falling in the same or similar categories are grouped and reassigned to a new annotated method, based on a custom conversion table (S2). A number of method codes are reassigned into an unspecified method category (reassigned annotation: UNSPM): these entries are logged and discarded. The user can select, at submission, whether they intend to use a lenient or a stringent method filter. The stringent filter will remove the interactions associated with the following codes: MI:0045; MI:0400; MI:0401; MI:0686; MI:0035; MI:0013; MI:0892; MI:0493; MI:0492; MI:0046; MI:MOV34-34KD; MI:HRIHFB2281; MI:0968; MI:0105; MI:bK984G1.4; MI:S2; MI:0063; MI:0110; MI:0058; MI:0101. While the lenient method filter will only remove interactions that are classified as: MI:0686; MI:MOV34-34KD; MI:0105; MI:bK984G1.4; MI:S2; MI:0063; MI:0110; MI:0058; MI:0101. The remainder will be classified as “general methods” and not removed.

Of note, the method conversion table converts the majority of the interaction detection method codes, particularly the most used ones. However, the conversion table is not complete by definition, since new method codes can be generated and may be annotated into primary database entries. To cope with this issue the default setting implemented in the automated tool is to ignore entries annotated with an unrecognized code. However, this setting can be modified when using the custom tool. In this case, users have the option of running the method reassignment function allowing the unrecognized methods to be retained. In the latter scenario, a record is generated in a temporary folder reporting the non-reassigned entries to be corrected manually.

#### *Data Processing: Interaction Confidence Score*

Entries are scored based on: i) the number of distinct methods that have been used to detect a specific PPI (method score, MS), and ii) the number of publications that report it (publication score, PS). A final score (FS) is calculated for each data entry by addition of the two previous scores (MS+PS=FS). The FS for each interaction is positively correlated with the confidence of that interaction (i.e. the greater the final score, the greater the confidence in the interaction).

### *Caenorhabditis elegans (taxonomy ID: 6239)*

#### *Query Submission, Data Acquisition and Data Processing and Scoring*

Upon submission, the internal C. elegans PPI network is queried to extract interactions where the seeds are classified as “interactor A” or “interactor B”. The list of interactions for each of the seeds is then scored based on the number of different interaction detection methods and PMID exactly as explained above for the human pipeline. The user is the provided with the scored list of PPIs and with a log file containing the list of proteins that were not present in the C. elegans PPI network (for which PPI informations are not available in PINOT). Of note, as for the human pipeline, the user can select, at submission, whether they intend to use a lenient or a stringent method filter. The internal C. elegans PPI network has been obtained by processing the Alliance_molecular_interactions.tar file downloaded from the Alliance of Genome Resources on 15th April 2019 (https://wormbase.org/#012-34-5). The processing consisted in the removal of all non protein-protein interactions, the removal of the unwanted fields keeping only: interactor.A; interactor.B; Alias.A; Alias.B; Interaction.detection.method; Publication.Identifier. We have converted the interactions detection method as previously described for the human pipeline.
